# Oviposition-related behaviours of *Limenitis camilla* in a common garden experiment

**DOI:** 10.1101/2023.02.06.527247

**Authors:** M. Marcantonio, R. Vodă, D. Da Re, Q Igot, R.L.H. Dennis, A. Vielfaure, S.O. Vanwambeke, C.M Nieberding

## Abstract

Human induced environmental changes are accelerating at an unprecedented pace, forcing organisms to rapidly adjust their behaviours. There is broad evidence that the main driver of the ongoing biodiversity crisis is land-use change, that reduces and fragments natural habitats. However, the consequence of habitat fragmentation on behavioural responses of fitness-related traits such as oviposition site selection in insects, which represent about 50% of \ Earth’s species diversity, have been so far understudied. In herbivorous insects, oviposition-related behaviours determine larval food access, and thus the fate of the next generation. We present a pilot study to assess differences in oviposition-related behaviours in *Limenitis camilla* butterflies from Wallonia (Belgium), one of the most fragmented regions in Europe. We first quantified variation in functional habitat connectivity for *L. camilla* across Wallonia and found that fragmented habitats had more abundant, but less evenly distributed *Lonicera periclymenum*, the host plant of *L. camilla*. Secondly, we compared in a semi-natural experimental setting the behaviours of field-caught *L. camilla* females originating from habitats with contrasted landscape connectivity. We found differences in behaviours related to flight investment: butterflies from fragmented woodlands spent more time in non-compass orientation flight, which we associated with dispersal, than butterflies from homogenous woodlands, where *L. periclymenum* was less abundant and more evenly distributed. Although results from this study should be interpreted with caution given the limited sample size, they provide valuable insights for the advancement of behavioral research that aims to assess the effects of global changes on insects.

## INTRODUCTION

Earth biodiversity has never been as rich as it is today, but at the same time, the rate of species extinction is higher than ever before due to human activities (Barnosky et al. 2011). A recent global assessment of biodiversity and ecosystem services has shown that a quarter of all known species are threatened, whereas about one million are already facing extinction in the short term (IPBES 2019). The current biodiversity crisis is mostly driven by landuse changes, which increase habitat fragmentation and loss and reduce the quality of remaining natural habitats (Díaz et al. 2019), resulting in cascading negative effects on the biota (Sih et al. 2011). Particularly susceptible to contemporary changes are insects, the most species-rich group of organisms on Earth (Bar-On et al. 2018). Insects, including many butterfly species, are vulnerable to human-induced rapid environmental changes (“HIREC” hereafter) and especially to land-use changes, including modern industrial (so-called “conventional”) farming and forestry practices (Warren et al. 2021). Butterflies are key bioindicators of habitat quality and flagship species for documenting the ongoing biodiversity crisis since their population trends and distribution changes have been uniquely monitored for decades (van Swaay et al. 2011). Currently, about 19% of all European butterfly species are in the IUCN categories threatened or near threatened, with populations rapidly declining (Van Swaay et al. 2011). Extinction risk of European butterflies is highly associated with habitat loss and fragmentation, in particular due to impacts on larval host plants or adult habitats (Van Dyck et al. 2009; Sánchez-Bayo and Wyckhuys 2019). During their lifetime, butterflies depend on multiple habitats, which taken together form their “functional habitat” (Dennis 2010) and the degradation of one or more components of their functional habitat imposes important constraints on their population dynamics. Butterflies that rely on few larval host plants and have stricter habitat requirements were shown to be more at risk of extinction than species with a larger range of preferences (Kotiaho et al. 2005), similar to findings in other insects (review: Gallagher et al. 2015; moths: Mangels et al. 2017; Habel et al. 2019).

HIREC has consequences for both the availability and configuration of suitable habitats and these changes may alter organism-functional habitat relationships through changes in behaviour (Wong and Candolin 2015). Behavioural changes affect in turn species interactions, population dynamics, evolutionary processes and biodiversity patterns (Sih et al. 2011; Candolin and Wong 2012). How animals respond to HIREC, and how this influences population survival, is an area of growing research interest (Bost et al. 2015; Prop et al. 2015; Teitelbaum et al. 2016). Yet, despite the vital links between environmental change, behaviour and population survival, the study of the interplay between behavioural phenotypes under HIREC is still at a very early stage. Behavioural change has the potential to mitigate or exacerbate the influence of environmental heterogeneity in the short time frame of HIREC (Candolin and Wong 2012; Gil et al. 2020). Behaviour change has thus emerged as a possible major evolutionary mechanism for species to face HIREC, both at the theoretical (Botero et al. 2015) and the empirical level (Sih et al. 2016; Dukas 2017; Snell-Rood et al. 2018; Ducatez et al. 2020).

One documented behavioural change related to abrupt and rapid alterations of the environment is linked to movement (Candolin and Wong 2012). Movement of an organism between different patches of its functional habitat relates to different life functions acting across multiple spatial and temporal scales, and the decision to move is based on the individual internal state, motion and navigation capacity and the external factors in the environment that affect its habitat (Nathan et al. 2008; Bestion et al. 2019). Behavioural responses to fragmentation related to movement can vary across taxa: different bird species increase their habitat by also using suboptimal areas (Siffczyk et al. 2003; Evens et al. 2018), some large mammals manifest avoidance of large portions of their critical habitat range and prefer staying in familiar areas, as a consequence of anthropogenic development (Northrup et al. 2015; Marchand et al. 2017). Insects were shown to avoid dispersal as well: *Bombus veteranus* bumblebees preferred to stay inside fragmented patches rather than visiting more distant areas, thus reducing pollen dispersal and increasing inbreeding of the visited plants (Goverde et al. 2002). Reduced mobility was also documented in butterfly populations originating from fragmented landscapes: *Papilio machaon* and *Maculinea arion* had reduced thoracic size, which may be linked to a reduced flight capacity (Dempster 1991). On the contrary, a heavier thorax that allowed better dispersal was found in isolated *Hesperia comma* populations (Hill et al. 1999). Land-use intensification was also shown to favour generalist over specialist-associated traits in butterfly communities, including increased dispersal and longer distance flight (Börschig et al. 2013). An increase of the costs related to dispersal was shown in the butterfly *Proclossiana eunomia* in fragmented landscapes; butterflies in fragmented landscapes displayed a more straight and longer flight (Schtickzelle et al. 2007).

Beside movements, changes in reproductive behaviours in butterflies are crucial when animals must deal with habitat reduction and fragmentation because of their consequences on the availability of their host plant. As habitats become increasingly limited and fragmented, female butterflies may allocate more resources for longer flights to locate the host plants and less resources for egg production, which may result in a resource allocation trade-off between egg production and flight investment (Gibbs and Van Dyck 2009). In herbivorous insects with limited mobility in the larval stage, such as butterflies, the oviposition site is also the offspring’s food source, making dispersal movements of butterflies for oviposition site selection decisive. Indeed, egg-laying choices in herbivorous insects have consequences on offspring growth (Gripenberg et al. 2010), defence (Denno et al. 1990) and competition (Anderson and Löfqvist 1996). A study on the butterfly *Maniola jurtina* estimated gene flow as an indirect method to account for reproductive success and found that dispersal is higher in landscapes similar in structure to this species optimal habitat (grasslands), as opposed to arable lands and woodlands, indicating that dispersal through unsuitable habitats (i.e. the “landscape matrix”) can reduce gene flow and thus reproductive success (Villemey et al. 2016). *Pararge aegeria* butterflies from fragmented, agricultural landscapes, where host plants may be distributed more widely, laid fewer but larger eggs in comparison with butterflies from woodland landscapes (Gibbs and Van Dyck 2009). Moreover *P. aegeria* from agricultural habitats displayed a more exploratory behaviour related to oviposition site selection (Braem et al. 2021a). Conversely, butterflies of other insect species may employ other strategies, produce more eggs of a smaller size and spend less time searching for high quality oviposition sites (Papaj 2000).

In this context, we addressed the question of whether habitat fragmentation has an effect on behavioural strategies related to oviposition site selection and other movements in the woodland specialist butterfly *Limenitis camilla* (Lepidoptera: Nymphalidae). We collected adult butterflies from woodland sites that have remained conserved and connected in the last five decades (i.e. “homogeneous habitats” hereafter), as well as from smaller, patchy woodland remnants (i.e. “fragmented habitats” hereafter) across Wallonia, Belgium. We quantified the activity of wild-caught butterflies in outdoor cross-shaped experimental cages, by focussing on flight activities in relation to site selection for oviposition, while controlling for the effect of abiotic factors such as temperature, humidity and insolation. Our overarching goal was to assess whether butterflies originating from homogenous vs. fragmented habitats showed behavioural differences related to movement and oviposition in a semi-natural outdoor setting. We answered the following specific questions:

- Does the level of spatial and temporal connectivity of *L. camilla* habitats vary across Wallonia? We expected (E1) that in this region there were areas with diverse habitat-change trajectories that could be pinpointed in space and time.
- Does host plant abundance and spatial distribution differ between homogenous vs. fragmented habitats? We expected (E2) that host plants were more abundant and more evenly distributed in homogenous compared to fragmented habitats.
- Do butterflies modify their behaviour due to the presence of the host plant in the experimental setup? We expected (E3) that butterflies spent relatively more time in the tunnel with the host plant than in the control tunnels, while accounting for abiotic factors that could have affected their movements in the tunnels.
- Do butterflies originating from fragmented vs. homogenous habitats differ in the time they allocate to different movement behaviours in the presence of host plants? We expected (E4) that individuals from fragmented habitats were more dispersive and spent less time navigating the tunnels in search of the host plant.

## MATERIAL AND METHODS

### The model species *Limenitis camilla*

*Limenitis camilla* is a widespread species in the Palearctic, ranging from temperate Europe to Turkey and extending as far east as Korea and Japan, and is currently present across all Belgium where it is strictly associated with remnant woodland habitats. This species is locally common in southern areas of Belgium, and since 2004 populations have been increasing in abundance also in the north of the country (De Prins and Steeman 2021). *Limenitis camilla* is usually univoltine, flying from mid or late June until mid-August and it occurs in moist broadleaved woodlands, where it is associated with open sunny areas in the tree vegetation cover (Bos et al. 2006; Dennis 2010). In Wallonia, *L. camilla* oviposit on *Lonicera periclymenum* (Dipsacales: Caprifoliaceae) and *L. xylosteum* (Dipsacales: Caprifoliaceae) (De Prins and Steeman 2021). *Lonicera periclymenum* is a vine, which prefers acidic terrain and grows close to the ground in the interior of temperate oak forests, whereas in canopy gaps, forest fringes and wooded banks it is a flowering winding climber (Grashof-Bokdam et al. 1998). *Lonicera xylosteum* is a small shrub that occurs on well-drained and calcareous soils in beechwoods or open mixed woods (Fox 1996). *Limenitis camilla* prefers plants located in semi-shade or dappled light inside the forest or at their edges (Fox 1996) and its larvae are not able to develop under full-sun conditions due to the sticky secretions of *Lonicera* leaf trichomes that hinder the construction of the larval hibernaculum (Fox 2005).

### Butterfly occurrence dataset

Butterfly occurrences for Wallonia were kindly provided by the Direction de la Nature et de l’Eau du Département de l’Etude du Milieu Naturel et Agricole de la Direction Générale Opérationnelle Agriculture, Ressources Naturelles et Environnement (DNE-DEMNA-SPW-ARNE et collaborators, agreement contract of sharing data n°CMDD1351). The database comprised 1,825 geo-referenced *L. camilla* observations spanning from 1970 to 2019. We aggregated *L. camilla* occurrences by using a 5 km hexagonal grid, considering the number of observations, the number of years with observations and the number of decades with observations (Fig. 1). This data aggregation step was pivotal to select areas in Wallonia where *L. camilla* populations are both currently abundant and stable in time.

**Fig. 1.**
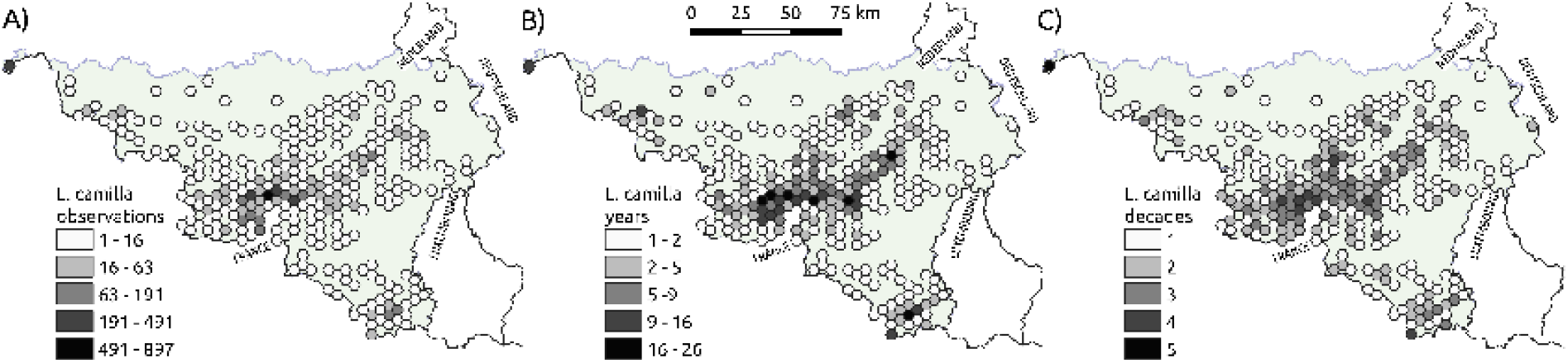
Maps with 5 km hexagonal grid reporting observations of *Limenitis camilla* in Wallonia from 1979 to 2019: A) the total number of observations; B) the number of years with at least one observations and C) the number of decades with at least one observation.

### Functional habitat connectivity and selection of sampling areas

#### Land use analysis

We selected *L. camilla* populations originating from broadleaved woodlands with different degrees of spatial connectivity and temporal stability in Wallonia. To pinpoint these forests we used standard, open access land cover datasets (Fig. 2): “LifeWatch-WB ecotope” (ECO) for the years 2006 and 2018 (http://maps.elie.ucl.ac.be/lifewatch/ecotopes.html), “ESA Corine Land Cover” (CLC) 1990 (https://land.copernicus.eu/pan-european/corine-land-cover/clc-1990/#) and “Les forêts anciennes de Wallonie” (Ancient Forests), representing the estimated forest coverage in Wallonia during the 18th century and based on the “Ferraris map” (https://opac.kbr.be/LIBRARY/doc/SYRACUSE/16992733). Each of these spatial layers was imported in the software GRASS GIS v7.9 (Neteler et al. 2012) and subset by selecting only land cover classes corresponding to broadleaved forests, preferred by *L. camilla*. This land use category corresponded to class 60 in ECO 2018, 2006 and CLC 1990 and to “boisement feuillu” and “forêt ancienne subnaturelle” in the Ancient Forests database. These four datasets were first transformed in binary raster data (1=forest, 0=matrix) with a spatial resolution of 10 m using bilinear interpolation and were then used to derive matrix-edge-core maps (MAC) using the open-access software LS Metrics (Niebuhr 2020). For this step, we defined an “edge” as the 100 m swath (or 10 pixels) area between the matrix and the core of a broadleaved forest patch (Matlack 1994). As a further step, we derived the distances between *L. camilla* observations and the edge of forest habitats (from ECO 2018) and compared it with all other butterfly observations in our dataset. A two sample t-test showed that *L. camilla* was found at signific-antly shorter (interior) distances from forest edges (mean=-34 m, SD=106) than all other butterflies observations (mean=51 m, SD=153 m; t(283)=12.4, p<0.001). We consolidated this information by using the interquartile range (IQ) of *L. camilla* distance-distribution (IQ range=-86,+11 m) in order to select an additional area around the 50 m forest edge which accounted for the ecology of both *L. camilla* and its host plants *Lonicera periclymenum and L. xylosteum*, while also considering the spatial distribution of recent butterfly observations. We defined this “extended forest edge” (EFE) swath as the functional habitat of *L. camilla*, or the habitat wherein the species is able to complete its life cycle – thus including mating, breeding and foraging habitats. We extracted pixels belonging to this category from each of the four spatial layers in sep-arate binary raster maps (EFE maps; sensu Dennis et al. 2006; Dennis 2010). These “functional habitat” maps were used as input in LS Metrics to derive functional connectivity (FC) maps, by considering a crossing capability threshold of 500 meters. This crossing capability can be assumed to approximate the maximum Euclidean distance that *L. camilla* butterflies fly from a functional habitat patch to reach another functional patch (see for example Dennis and Shreeve 1996). In addition, this figure is in the range of the dispersal capability of larger and dispersive butterfly species (e.g. *Gonepteryx rhamni*, whose dispersal ability between habitat patches is > 1 km) and smaller and more sedentary species (e.g. *Leptidea sinapis*; 300 m). The FC maps had thus pixel values corresponding to the functional habitat area in hectares available for a putative dispersing *L. camilla* located in that pixel. We derived two summary FC maps reporting the temporal average and coefficient of variation by considering the four FC maps. As a final step, these two maps were aggregated using a hexagonal 1 km grid which allowed us to obtain spatial units that could be readily used as reference sampling locations. Thus, the final 1 km maps represented the average functional habitat connectivity and its variability over the four considered time points (18^th^ century, 1990, 2006 and 2018).

**Fig. 2.**
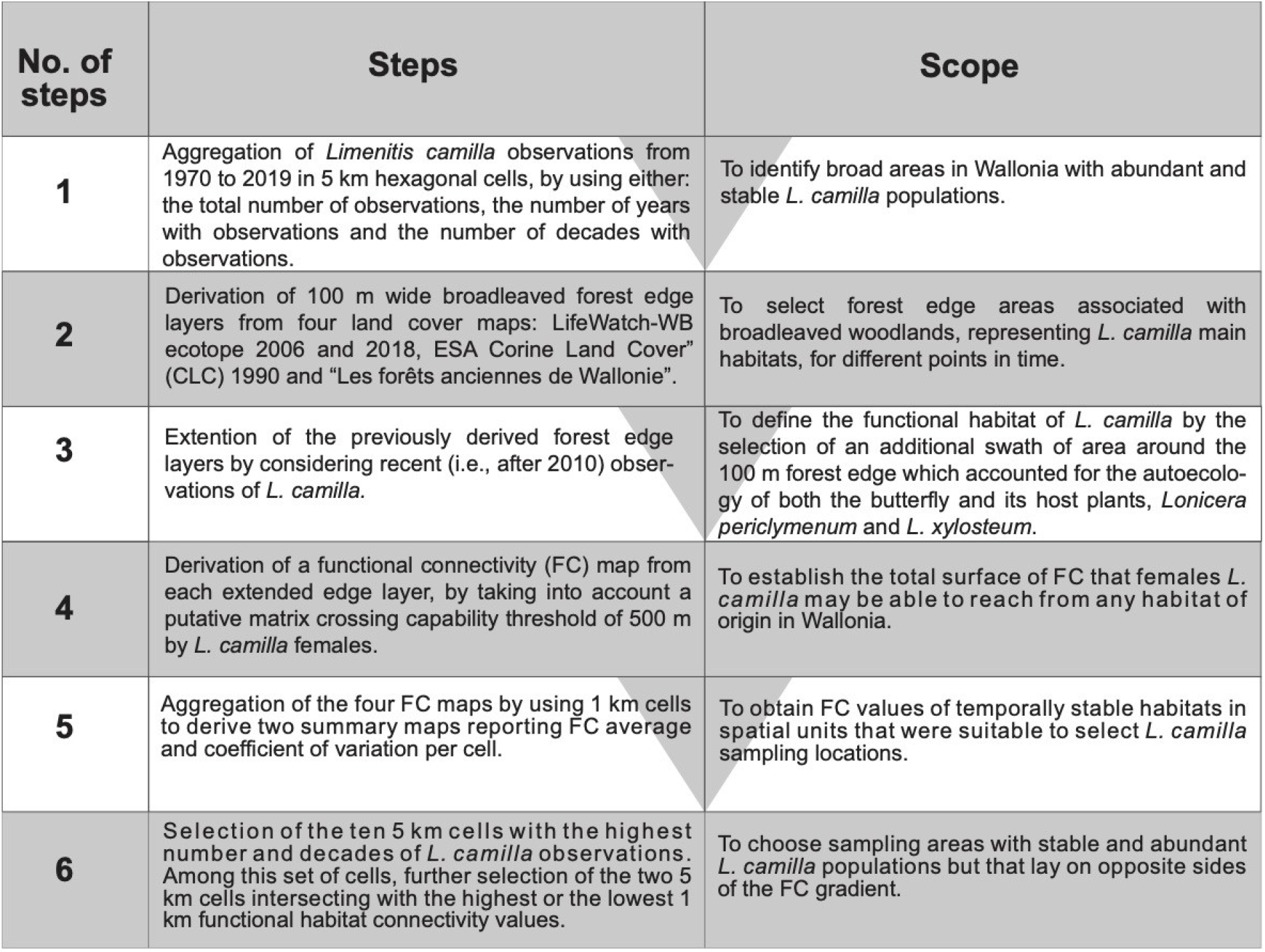
Steps followed to define the functional habitat for *L. camilla* and to choose the collection sites with stable and abundant *L. camilla* populations in Wallonia.

#### Sampling populations of *L. camilla* and its host plant *L. periclymenum*

We sorted the 5 km hexagonal cells (see Fig. 1) by using the highest number of occurrences, and then decades and years with observations for Wallonia as partial sorting criteria. Afterwards, we selected two 5 km cells with respectively the highest and lowest 1 km functional habitat connectivity values. These 1 km cells represented the areas where we collected *L. camilla* (from the end of June to the beginning of August, thus we assumed that all butterflies were collected after mating) and surveyed for the presence and abundance of its host plants. We chose to assess presence and abundance of the host plant inside or in the vicinity of the same 1 km hexagonal cell where *L. camilla* were also sampled. Thus, we selected only EFE habitats inside each 1 km cell where butterflies were collected and drew ten random points in-side each of these areas using QGIS v3.16 tool “Random points inside polygons”. For the random point draws we added a constraint of a minimum 200 m distance between points to avoid pseudo-replication due to spatial autocorrelation. A subset of these points was used as central coordinates to build 20×20 m vegetation surveys aligned along cardinal directions. We divided the plot in 5×5 m quadrants, where we recorded the presence and total number of host plants stems (limiting the count to a maximum of 25 stems per plot due to time constraints), in order to have more resolution for assessing host plant spatial distribution.

### Behavioural experiments and data analyses

Upon field collection, butterflies were kept in individual 11×11×14 cm transparent plastic boxes placed inside Sanyo incubators under constant conditions (temperature day/night: 22°C/16°C; relative humidity: 60%; photo-period light/dark: 2h/22h under solar light spectrum simulating lamps; Philips HPI-T Plus 400W/645) for at least 24 hours before behavioural experiments. These conditions were intended to reduce activity and thus wing wear, whereas stimulating activity and oviposition during behavioural trials. All butterflies had access to artificial nectar (20% sugar solution) and water ad libitum during this phase.

We conducted behavioural experiments in outdoor flight arenas, which consisted of four cross-shaped greenhouse aluminium tunnels, each 8×3×2 m covered with a thin black insect mesh and placed in the university experimental forest “Bois de Lauzelle” (part of Natura 2000 site “Vallée de la Dyle à Ottignies”; Ottignies, Wallonia; 50.680434 N, 4.614885 E; Fig. 3). *Limenitis camilla* and its host plants *L. periclymenum* are both found in Bois de Lauzelle. Healthy and fully developed *L. periclymenum* (*L. xylosteum* was not considered since it was not found in any sampling areas) used as host plant during behavioural experiments were collected nearby the flight arenas, re-planted in a textile pot, and placed in one of the four tunnels of the experimental cage defined as the “target tunnel”. Whereas the other three arms were left empty as “control tunnels”. The identities of target and control tunnels were kept constant across repeated trials for each butterfly. A temperature and humidity sensor (HOBO U23 Pro v2 Temperature/Relative Humidity Data Logger, Onset, Massachusetts, USA) was attached to the ceiling at the end of each tunnel to acquire data on microclimatic variation between tunnels (recorded every 5 minutes for the whole study duration).

**Fig. 3.**
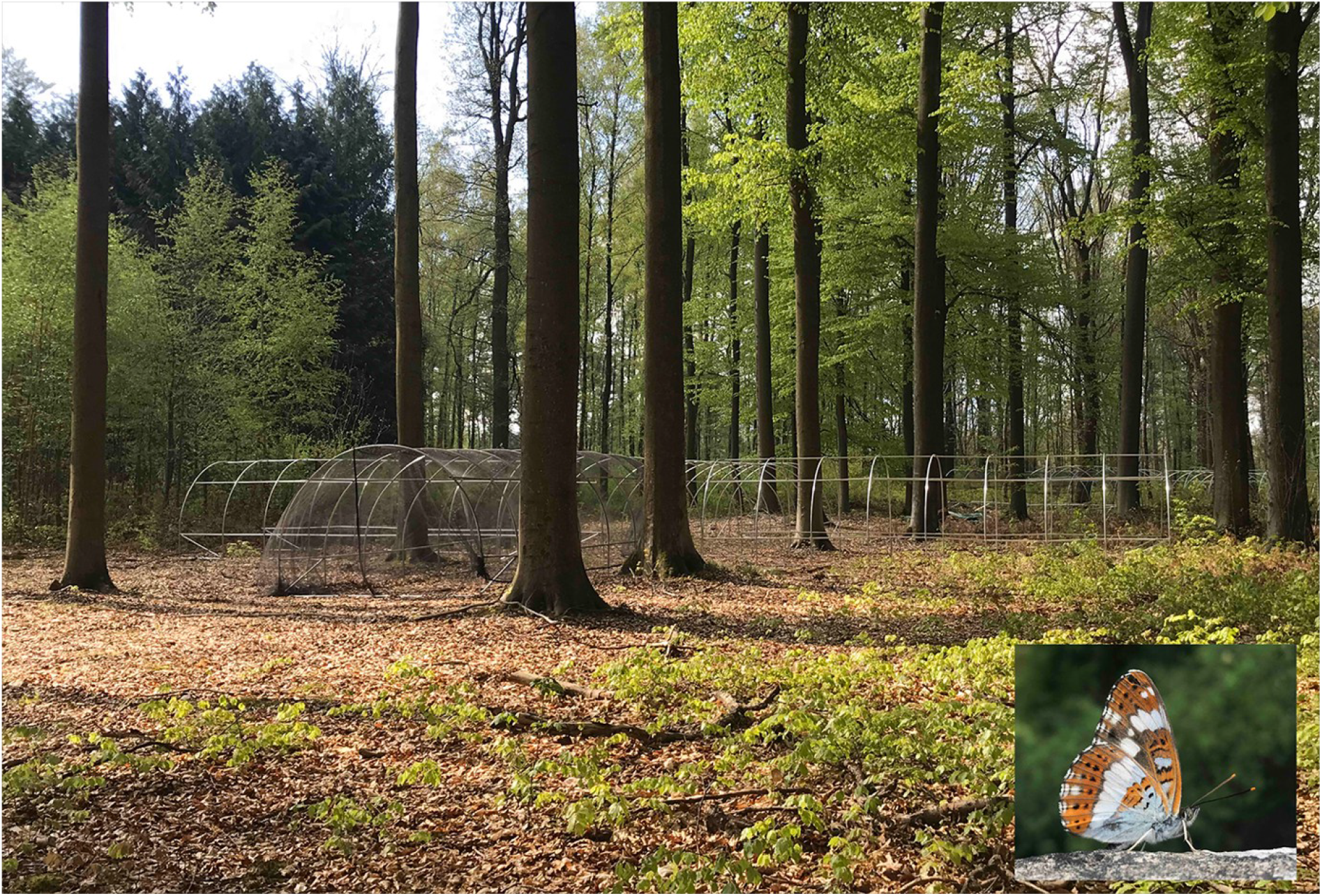
An outdoor flight setup consisting of four cross-shaped greenhouse aluminium tunnels covered with insect mesh, placed in “Bois de Lauzelle” forest, an experimental site belonging to UCLouvain, and an inset representing the study species *Limenitis camilla*.

Before behavioural trials, butterflies were sensitised to lower their threshold to oviposition behaviour during following behavioural trials. Butterflies were released and presented with the host plant for 10 minutes in a 2×3×2 m temporary enclosure built at the end of one of the flight tunnels. Butterflies that did not oviposit during a sensitisation trial were tested again the following day, whereas butterflies which oviposited were considered “sensitised” and used later the same day for behavioural trials.

Behavioural trials were carried out in the whole experimental arena (Fig. 3). Butterflies were brought to the central part of the arena inside plastic boxes whose cover was loosened to be removed with a long stick from an operator outside the arena. A behavioural trial started when the individual flew out from the box and lasted for 30 minutes; during this time butterfly behaviours and movements were voice-recorded. Trials were stopped immediately upon oviposition. Leaves on which butterflies oviposited were removed to avoid biased behaviour in successive trials as eggs are known to be used as social cues by some butterfly species. Behavioural trials were conducted on each butterfly over successive days, until death or when wing wear did not allow flight. Butterflies were not provided with a host plant for oviposition between behavioural trials in order to increase motivation for egg laying in the behavioural experiments.

During experimental trials we monitored and voice-recorded all butterfly behaviours, which were subsequently pooled into three behavioural categories by adapting to our model butterfly species the terminology proposed by Grob et al. 2021:

- “Non-compass orientation”: flight activities associated with either butterflies repeatedly flying into the covering net of the tunnels or with flight towards the sun, a known tendency in butterflies. We interpreted these behaviors as the initiation of a dispersal movement from one patch of potential habitat (in our case a patch inside the flight structure) to another. Indeed, butterflies have been found to fly straighter (possibly using non-compass orientation, such as wind or phototaxis) when dispersing in the landscape matrix than when flying inside habitat patches (Schtickzelle 2007). We interpreted an increased time spent in non-compass flight (e.g., phototaxis) in butterflies as indication of an increased investment in dispersal behavior.
- “Navigation”: all behaviors during which the butterflies interacted with the habitat resources inside the flight tunnels: flying or walking on the tunnel structure, on the ground and freely in the space inside the tunnels, flying over/around the host plant, landing on the host plant. Indeed, butterflies mostly use a directional type of flight behavior within habitat patches, based on landmarks or flowers for example, while searching for foraging or oviposition resources. This pattern has also been found in other organisms (Haughland & Larsen, 2004; Zavorka et al. 2015; Jablonszky et al. 2020).
- “Resting”: sitting still on the net, tunnel structure or on the ground;

We aggregated the durations (in seconds) of each behaviour category per individual, tunnel and experimental trial. Each behaviour was associated with tunnel-location of the host plant, average temperature, humidity and insolation, which were recorded during the experiments and averaged over each 30 minutes trial. This dataset was used to test whether there were differences in the total duration of the various behavioural categories among butterflies, by using Generalised Linear Mixed Models (GLMMs) with a negative-binomial link function due to the overdispersed distribution of duration data. We considered trial and butterfly ID as nested random factors whereas behavioural category, temperature, humidity and insolation as covariates. We next tested whether the presence of the host plant in a tunnel was associated with a different residence time in the tunnels. We further aggregated the dataset using the presence/absence of host plant in the tunnels as a grouping factor, thus discarding tunnel IDs. This dataset was used to model the duration of behavioural categories by adding an interaction term between target (with host plant) and control (without host plant) tunnels and behavioural category. Temperature, humidity and insolation were used as additional covariates. An offset term was added to this latter model to account for control tunnels having three-time higher probability of hosting behavioural activity with respect to the target tunnel.

The non-aggregated dataset was used to fit a third model to test potential difference in behaviours between butterflies with different origins, i.e., fragmented vs. homogenous woodlands. This model included an interaction term between behavioural category and origin of the butterfly. We visually inspected models’ residuals and used the Akaike Information Criterion (AIC) to select the final models. All the analyses were carried out in R 3.6.3 (R Core Team 2020) through the functions available in lme4 package (Bates et al. 2014). Both dataset and R code to reproduce all models are available at https://osf.io/2ej7x/.

## RESULTS

### Selection of habitats with low and high levels of functional connectivity within areas of frequent *L. camilla* observations (Expectation 1, E1)

The number of aggregated *L. camilla* observations per 5 km cell (henceforth abundance) ranged from 1 to 897 (Fig. 1). The highest abundance was found in south-west Wallonia (Namur province) and in the detached Comines-Warneton municipality (Hainaut province). The north, north-east and south-east parts of Wallonia lacked almost entirely observations for *L. camilla* in the past 40 years. The maximum number of years and decades with observations for a cell were respectively 26 and 5, with a spatial distribution similar to that shown by *L. camilla* abundance data (Fig. 1).

Stable functional woodland habitats were mainly distributed along a diagonal line stretching from the south-east of Namur province to the north-east of Liege province (Fig. 4). The connectivity analysis showed that regions with abundant highly connected habitats underwent only moderate changes when compared to areas with lower habitat connectivity, such as north-west Hainaut, west Wallon Brabant and south-east Liege (Fig. 4). These latter areas were indeed characterised by a higher temporal variability (e.g., loss) of functional woodland habitat cover and connectivity. Further we selected the two 5 km cells (see Fig. 1) having the lowest and highest average connectivity values respectively, among cells with the highest *L. camilla* abundance and persistence over the last five decades. These cells represented reference sampling locations in fragmented and homogenous woodlands.

**Fig. 4.**
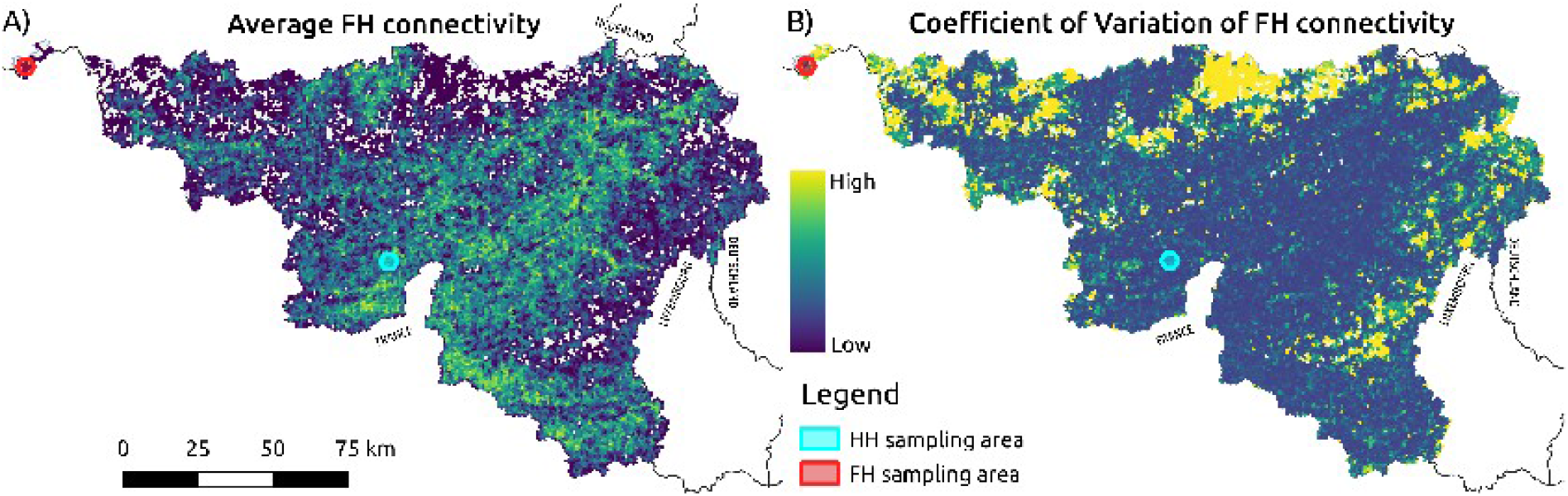
Maps reporting: A) the average functional habitat (FH) connectivity and B) its coefficient of variation (right) for Wallonia. Transparent cells inside Wallonia do not contain FH. The red and light-blue small hexagons overlaid on the maps show homogenous (HW) and fragmented (FW) woodlands from where butterflies were collected.

The cell with the lowest connectivity values (i.e., area of functional habitat available to a butterfly that is able to fly 500 m over the landscape matrix) was located in the detached enclave of Comines-Warneton (red hexagon in Fig. 4). This area was characterised by very low functional habitat connectivity (average 0.2 km^2), while showing moderate *L. camilla* abundance: 342 individuals observed in 11 different years in the last five decades. We further verified that the low connectivity of this detached enclave was not due to an “edge effect” by using land cover additional datasets for both Flanders and France (see SI1 for further information). The cell with the highest connectivity (average 91 km^2) values was centred on the village of Doische. This cell also contained a high number of *L. camilla* observations, 491, collected in 21 years over three decades (light-blue hexagon in Fig. 5). Inside these two 5 km cells, we selected the 1 km cells with the highest cover of forest functional habitat (Fig. 5). Inside and in the proximity of these 1 km areas we collected *L. camilla* and assessed the abundance of their host plant species.

**Fig. 5.**
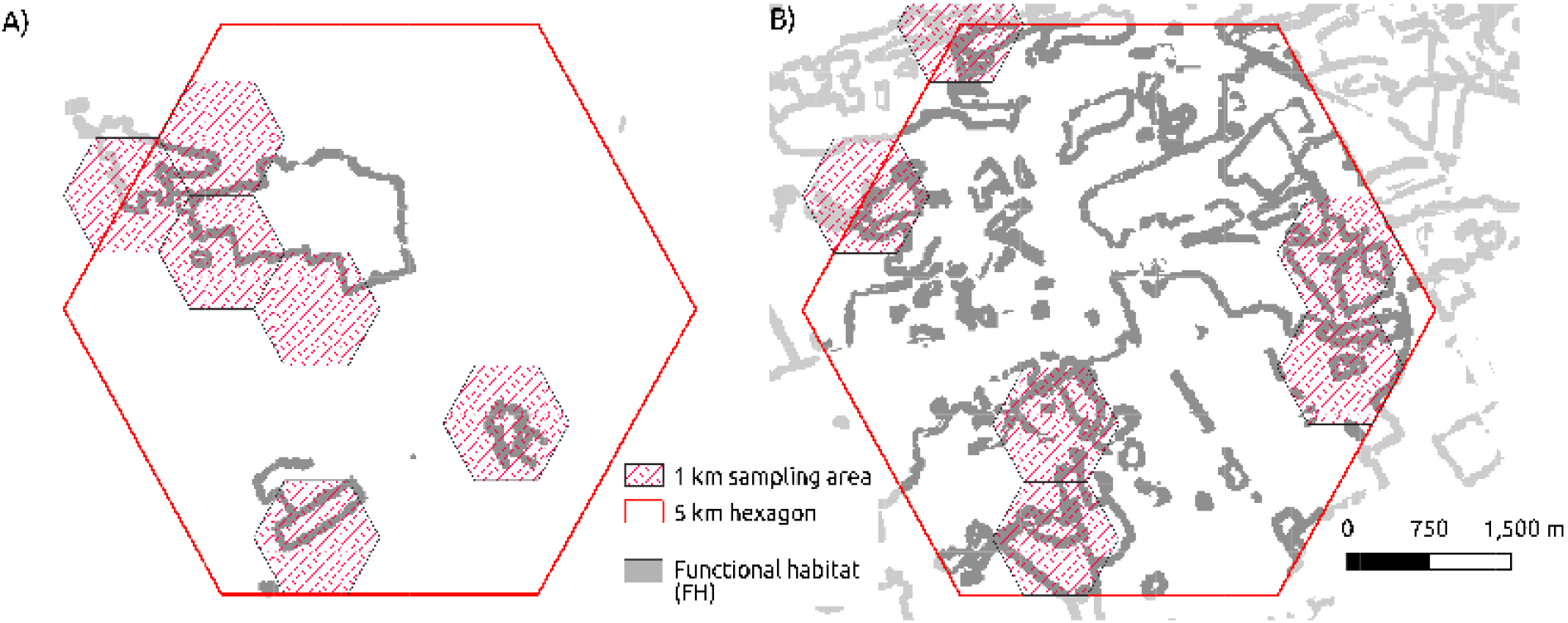
Sampling areas: A) Comine-Warneton cells (fragmented woodlands) and B) Doische (homogeneous woodlands). The red cells with canvas pattern were 1 km cells chosen for collecting *L. camilla* and for quantifying host plant abundance. The background map reports in gray the extent of functional habitat.

### Spatial distribution and abundance of the host plant (Expectation 2, E2)

We carried out a total of 10 host plant surveys, 6 in homogenous woodlands and 4 in fragmented ones and found *L. periclymenum* in 8 out of the 10 total surveys, while *L. xylosteum* was not found in any areas. We fitted a Linear Mixed Model (LMM) considering plot ID as a random factor to test for differences in the number of *L. periclymenum* stems between the two areas. Model results showed that the sampled areas in fragmented habitats had an overall higher abundance of the host plants with respect to homogenous habitat areas (LMM; *β=*6.25, *df*=34.45, *p*=0.015). On the contrary, the Shannon index calculated at survey quadrant level was higher in homogenous habitats (H=2.23, sd=0.19) suggesting that *L. periclymenum* was more evenly distributed in these areas than in fragmented habitats (H=1.83, sd=0.28).

### Behavioural activity of populations from fragmented vs. homogenous woodlands (Eexpectations 3 and 4, E3 and E4)

We collected a total of 46 *L. camilla* (from 29/06/2021 to 11/08/2021), 40 from homogenous and 6 from fragmented woodlands, and 36 underwent one to six 10-minutes sensitisation tests. During a total 71 sensitisation tests, 10 butterflies laid eggs, 7 from homogenous and 3 from fragmented habitats (representing 17% and 50% of the two groups respectively), during 11 different tests. Twenty-six 30-minutes behavioural trials were carried out on 10 butterflies from homogenous and 4 from fragmented woodlands, which oviposited during sensitisation or underwent at least 3 sensitisation tests. Only one *L. camilla* (from a homogenous woodland) oviposited (twice) during behavioural trials. Butterflies spent more time resting (R=90.9%, 1.1% of which was represented by walking) compared to non-compass orientation and navigation behaviours, which represented 5.1 and 4.0% of the total recording time, respectively (Fig. 6). We further assessed linear combinations of model covariates, which revealed how resting durations were significantly higher than both non-compass orientation and navigation flights, whereas time spent in these latter categories did not differ between them (μ=-0.552, p-value=0.66). In addition, butterflies spent significantly more time resting, and tended to exhibit non-compass orientation less and to navigate more, in the target tunnel with respect to the other tunnels (Table 1).

**Table 1.** Summary table for a model testing time spent by butterflies in the target tunnel (TT, i.e. tunnel with the host plant) in the three different behavioural categories, by also considering the interaction between tunnel type and behavioural category. Random factors: σ trial=3.7e-06; σ id=6.5e-07. NCO=non-compass orientation, N=navigation.

| Model term | Estimate | Std. Error | z value | Pr(>\|z\|) |
| --- | --- | --- | --- | --- |
| Intercept | 3.62 | 0.21 | 17.46 | <0.001 |
| Tunnel TT | 2.05 | 0.36 | 5.94 | <0.001 |
| *Non-compass orientation (NCO)* | -2.58 | 0.33 | -8.32 | <0.001 |
| *Navigation (N)* | -2.61 | 0.41 | -5.85 | <0.001 |
| *TT: NCO* | -0.12 | 0.55 | -0.43 | 0.831 |
| *TT: N* | 0.74 | 0.67 | 0.71 | 0.262 |

**Fig. 6.**
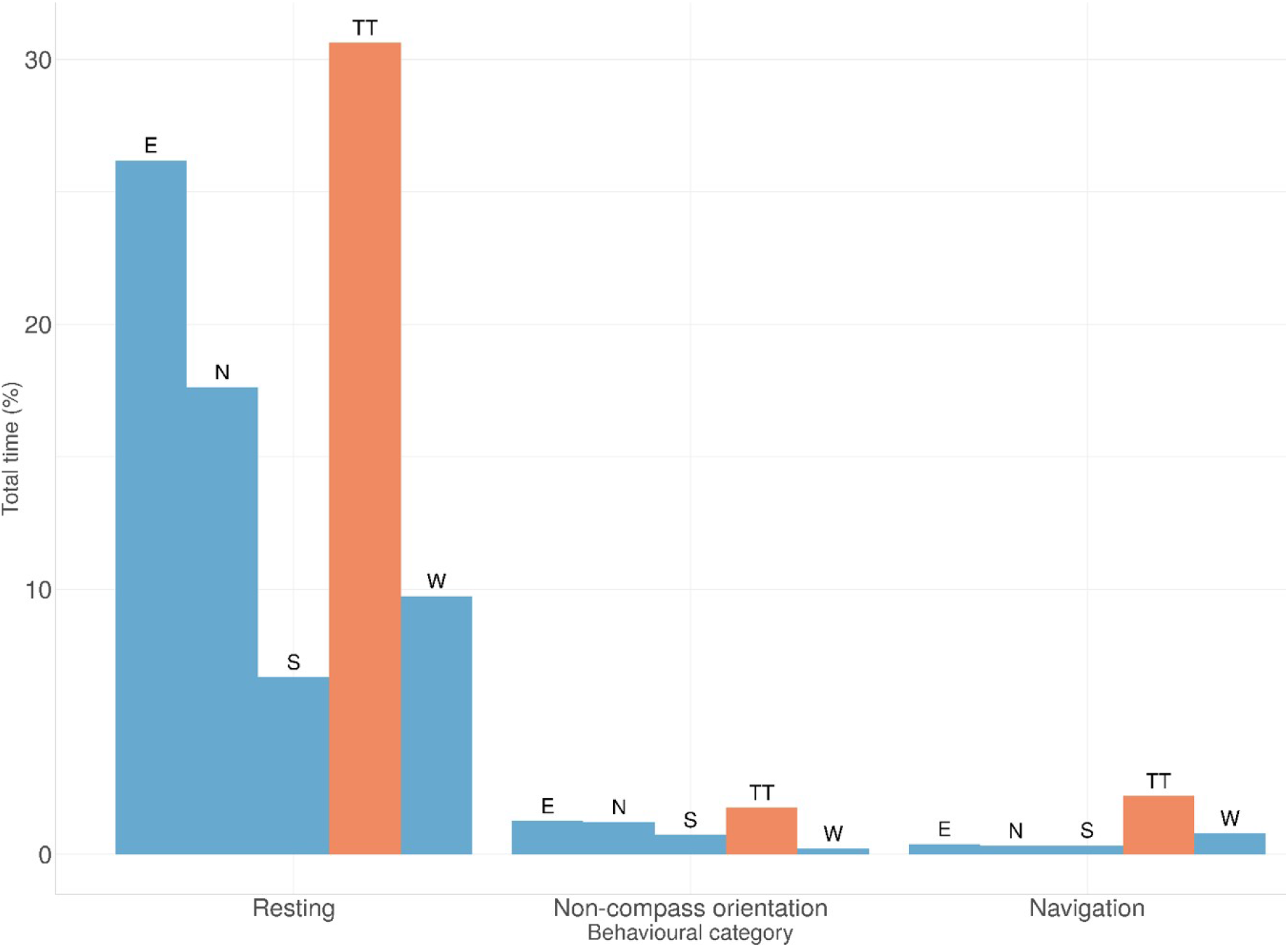
Barplots showing the percentage of total experimental time (for all 14 tested butterflies) spent in the different behavioural categories. Red bars indicate time spent in the target tunnel (TT), whereas blue bars in control tunnels (which can be any of the four tunnels named following their long-axis cardinal direction).

Overall butterflies spent considerably more time in the east and south tunnels (212 and 177 minutes respectively), which recorded higher average temperature during trials (25.5 and 23.9 °C respectively) than in north and west tunnels (113 and 60 minutes respectively), which recorded lower average temperatures (23.3 and 22.4 °C). Despite these results, temperature, humidity and insolation showed all very small, non-significant, model coefficients and did not change model AICs, thus were removed from all final models.

The origin of the tested butterflies did not affect the time they spent resting. On the contrary, the interaction term suggested that butterflies originating from fragmented woodlands spent more time in non-compass orientation flight with respect to butterflies that originated from homogeneous habitats. Navigation activities did not vary between habitats of origin (Table 2, Fig. 7).

**Table 2.** Modelled time spent in the three different behavioural categories according to the origin of the butterfly (HW=homogeneous woodlands, FW=fragmented woodlands; NCO=non-compass orientation, N=navigation). Summary table with random factors: σ trial=1.51e-06; σ id=6.8e-07, σ tunnel=3.03e-07.

| Model term | Estimate | Std. Error | z value | Pr(>\|z\|) |
| --- | --- | --- | --- | --- |
| *Intercept* | 6.76 | 0.22 | 31.26 | < 0.001 |
| *Origin FW* | -0.37 | 0.36 | -1.02 | 0.311 |
| *Non-compass orientation (NCO)* | -3.00 | 0.32 | -9.47 | < 0.001 |
| *Navigation (N)* | -2.61 | 0.40 | -6.57 | < 0.001 |
| *FW:NCO* | 1.16 | 0.58 | 1.98 | 0.043 |
| *FW:N* | 0.08 | 0.58 | 0.14 | 0.892 |

**Fig. 7.**
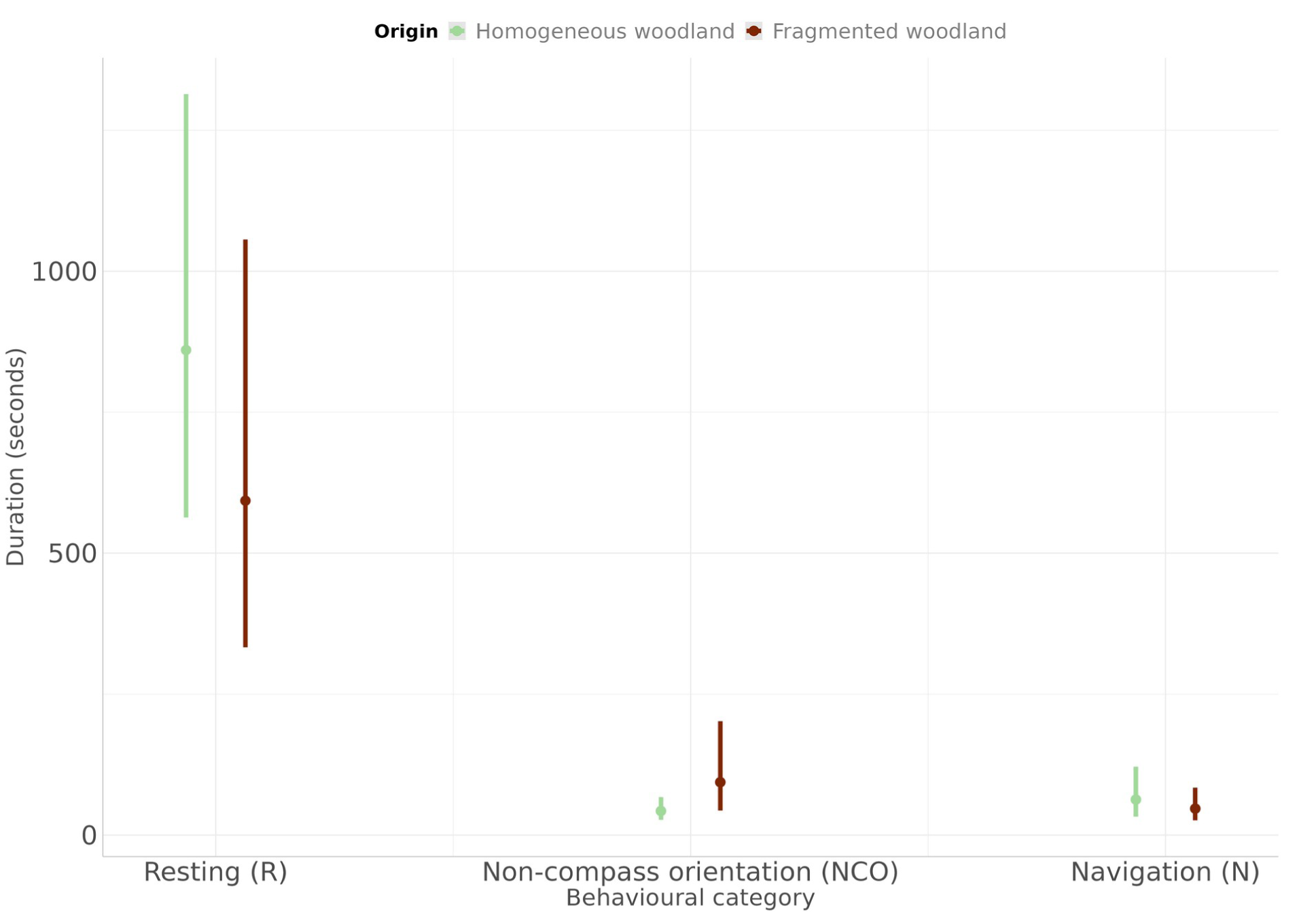
Variation in three main behavioural categories (resting, non-compass orientation and navigation) for homogeneous (in green), and fragmented (in brown) woodlands, with model marginal means and 95 % CIs from Table 3. The 95% CIs have been calculated considering only model regression parameters (i.e. not considering the uncertainty in the model variance parameters).

## DISCUSSION

Habitat fragmentation is one of the facets of the pervasive changes that are transforming Earth ecosystems and causing the ongoing biodiversity crisis (Ewers and Didham 2005). Species depending on specific habitats or resources are more affected by fragmentation due to their lower competition capacity in more isolated and reduced habitats (Chetcuti et al. 2020). Among the most intriguing and less investigated consequences of habitat fragmentation on biota are species behavioural responses whose phenotypic plasticity may impact their evolutionary trajectories (Candolin and Wong 2012; Nieberding et al. 2018; Snell-Rood and Ehlman 2021).

Here we conducted a pilot study in which we addressed the question whether habitat fragmentation has an effect on behavioural strategies related to oviposition in populations of the woodland specialist butterfly *L. camilla*. We first discuss our results on the connectivity of the functional habitat of *L. camilla* in Wallonia (E1 and E2) and then delve into comparing behavioural strategies of populations originating from homogenous and fragmented woodlands (E3 and E4). We discussed the limitations of this pilot study in a final paragraph, where we advise caution in the interpretation of our results due to the limited sample size while suggesting future ways forwards to overcome them.

### Functional habitat connectivity and host plant spatial distribution (E1 and E2)

We found areas in Wallonia with opposite trends in the connectivity of the functional habitat of *L. camilla*, represented by secondary forests characterised by open sunny areas in the tree vegetation cover (Dennis 2010; Vries et al. 2021). Western Europe suffered massive forest loss starting from the late prehistory and antiquity, followed by a reforestation phase and renewed deforestation in medieval times, with a dramatic acceleration during modern times (Woodbridge et al. 2015; Fuchs et al. 2015; Fyfe et al. 2015). These long-lasting land-cover changes often caused extreme fragmentation of remnant forest habitats; but areas with a more complex topography remained less exploited due to higher costs of land transformation (Ustaoglu and Collier 2018). South-western areas of Wallonia were characterised by larger “conserved” woodlands interrupted by smaller agricultural lands (homogenous habitats); here we found that the distance between woodland patches was often well into the average dispersal capacity of *L. camilla*, and thus these woodlands formed an extended network of connected functional habitats. In addition, in homogenous woodlands, *L. camilla* host plants were less abundant but more evenly distributed in space, producing less isolated and more regularly spaced oviposition habitats. In contrast, the fragmented woodlands were mainly present in the (less topographically complex) north and east boundary portions of Wallonia, where host plants were overall more abundant but also more isolated (i.e., aggregated). These differences in host plant spatial distribution in our study area may be explained by the autoecology of *L. periclymenum*, which prefers shady edge habitats where roots stay in the shade and shoots can climb up more easily to the sun. In Wallonia, fragmented patches of forest often presented sharp edges shared with agricultural or urban lands where *L. periclymenum* could become dominant. On the contrary, more extended and connected woodlands were characterised by frequent but less sharp woodland edges, which may explain an even spatial distribution of *L. periclymenum* in these areas (Grashof-Bokdam et al. 1998).

### Behavioural strategies of populations from fragmented vs. homogenous woodlands (E3 and E4)

Behavioural trials showed that *L. camilla* females remained overall more time in tunnels with host plants, regardless of the habitat of origin (confirming hypothesis E3). In addition, individuals from fragmented and isolated habitats spent more time in non-compass orientation with respect to individuals originating from connected functional habitats (confirming E4). Plastic behavioural responses related to flight and specifically oviposition site selection are fundamental for specialist butterflies that experience environments where their host plants (being somewhat limited) are more likely to be infrequent (coarse-grained environmental variation, Snell-Rood and Ehlman 2021). The observed increased investment in non-compass orientation flight may be therefore due to a positive selective pressure in butterfly populations from more isolated habitats. When faced with an environment where oviposition resources are limited to isolated small patches of functional habitat, butterflies may readapt to local conditions, for example exploiting a sub-optimal host plant, or they may disperse in search of more suitable habitats (see Singer 2015). A stronger investment in non-compass orientation flights, which for example may result in movements from one potential habitat patch to another, observed during the behavioural trials, may be a response to the perception of a non-suitable habitat in order to boost the likelihood of finding a new (disconnected) suitable habitat. Hence, we interpreted an increased time spent in non-compass orientation flight in butterflies from fragmented woodlands as an indication of an increased investment in dispersal behavior. However, our experimental settings prevented us to determine the ending point of non-compass orientation flights. Thus, we cannot rule out that non-compass orientation flight was associated with a more efficient strategy to exploit infrequent resources inside habitat patches and, as such, not necessarily associated with dispersal towards a different habitat patch. In line with this conjecture, we found that host plants were both more abundant and more aggregated in space in fragmented woodlands, producing more disconnected oviposition habitats than in homogenous woodlands. This finding may further suggest that the higher investment of butterfly populations from fragmented habitats in non-compass orientation may be adaptive, as it may help them find more isolated, but locally more abundant clusters of host plants in the same woodland patch (i.e., increasing the range of local environments experienced; Snell-Rood and Ehlman 2021). In this regard, a study on the effect of plant spacing on the movement of flea beetles found that less evenly distributed plant patches were associated with more random foraging patterns than when plants were more evenly distributed in space (Kareiva 1982).

Our study may thus provide insights on the potential selective pressure acting on butterfly populations facing intense habitat fragmentation and isolation. Increased investment in dispersal movements may reduce realised fecundity as additional energy is allocated to flying instead of oviposition (Gibbs et al. 2010; Saastamoinen et al. 2010; Bonte et al. 2012; Singer 2015). Previous studies found that highly mobile *Drosophila* genotypes showed lower long-term memory (Mery et al. 2007; Snell-Rood and Steck 2019), suggesting that movement behaviours and memorisation may be inversely selected for finding new suitable habitats. Moreover, it has been proposed that non-compass orientation (which could be related to the initiation of some dispersal behaviours) in insects may not require higher brain processes, whereas navigation behaviours involve the integration of positional information in the mushroom body (Grob et al. 2021; Franzke et al. 2020). Therefore, the higher duration of non-compass orientation observed in *L. camilla* individuals originating from fragmented habitats may be also correlated with a lower investment in higher brain centres, resulting in lesser spatial learning abilities.

Our findings add to a body of work on butterfly behaviour and conservation showing that butterflies occupying small, fragmented habitat patches display phenotypic changes, including changed mass allocation to flight muscles and to reproduction (Hill et al. 1999; Dempster 1991), compared to populations occupying more contiguous patches. Findings did not support the navigation assumption of hypotheses E4, despite the non-significant increased time spent “navigating” by butterflies from connected habitats (see Fig. 7). This result may be related to the short time that butterflies spent “navigating” during the behavioural trials, possibly due to the experimental conditions, as discussed in the following section.

### Limitations and perspectives

We used a large experimental setup placed in a habitat specific to *L. camilla* that provided ecologically relevant conditions to tackle behavioural strategies related to oviposition in butterflies (Nieberding et al. 2018). This approach allowed us to study *L. camilla* behaviours in conditions as close as possible to their natural environment. Such experiments cannot indeed be easily carried out for highly vagile species under full field conditions (but see Söderström and Hedblom 2007) and tracking devices are not yet light enough for experiments with European butterflies (Srygley and Kingsolver 2000; Daniel Kissling et al. 2014; Kaláb et al. 2021). We also faced practical complications that limited the sample size of tested butterflies. We were able to test only a small number of individuals (14 total out of the 46 collected) due to colder and wetter than usual weather conditions (for more information see SI2). Moreover, we tested butterflies in an outdoor arena, which, despite being much larger than common laboratory behavioural cages, may have not been optimal for wild butterflies. First, we conceived our setup as a cross-shaped structure that had the advantage of mimicking behavioural bioassays of four-way olfactometers used to study animal behaviour in laboratory conditions (Lin et al. 2016), and that had been also previously tested on another butterfly species (Braem et al. 2021). This type of bioassay is very flexible and allows for experiment reproducibility, but it has a low volume-perimeter ratio and thus a higher chance that tested individuals contact the covering net, perceiving it as a threat or impediment, which may not be suited for highly vagile flying species. Butterflies bumping into the covering net may have confused it for features of vegetation and hence tried to pass through it, which may explain the longer time spent in non-compass orientation flight than in activities related to spatial navigation. We observed that many butterflies, after initially contacting the net during a non-compass orientation flight, seldom changed their behaviour into navigation and usually continued to bump into the net or stopped altogether. Second, our experimental setup was placed in the interior of a beech forest, to avoid overheating and inactivity of the tested butterflies in case of extreme temperatures. This environmental setting was not optimal for behavioural trials as conditions were often too shady and cold for *L. camilla*, which may explain the long time they spent resting and the almost total lack of oviposition. Third, the arena was voided of vegetation or other habitat features to maximise the saliency of the stimulus provided by host plants in the arena, while avoiding other confounding stimuli, such as the presence of non-host or nectar plants. However, an unexpected side effect of this “simplified” environment might have been deprival of information on habitat quality for butterflies. Theoretical studies on species dispersal showed that individuals tend to emigrate from their current habitat patch more frequently when they do not have information on the surrounding habitat or on local population densities (Enfjäll and Leimar 2009).

Building upon the results and experience gained from this pilot dataset, we are developing a larger squared-shaped experimental arena with a higher ceiling, placed in an open field and with interior conditions richer in cues (e.g., social cues). We expect this setting to be better suited to study oviposition-related behavioural strategies in *L. camilla* and other butterfly species (see Saastamoinen 2007; Duplouy et al. 2013).

## ACKNOWLEDGMENTS

This research was funded by UCLouvain, the Fédération Wallonie-Bruxelles, and by the Fonds National de Recherche Scientifique (F.R.S.-FNRS). R.V. and M.M. were supported by the F.R.S.-FNRS grant number T.0169.21 to C.M.N.; M.M. was supported by the “Action de Recherche concertée” grant number 17/22-086 to C.M.N. We thank the ADPI and GPEX services (teams of A. Morise and T. Thyrion) at UCLouvain for providing technical support in setting-up the flight tunnels.

## STATEMENTS AND DECLARATIONS

### Author’s Contribution

C.M.N. had the idea of the study and C.M.N, M.M. and R.V. contributed to the design of the study. Material preparation and data collection were performed by M.M., A.V. and R.V. The spatial analysis was performed by M.M, D.D.R, Q.I and S.O.V., while the analysis of the behavioural data was performed by M.M. The first draft of the manuscript was written by M.M. and R.V. and all authors commented on previous versions of the manuscript. All authors read and approved the final manuscript.

